# Incorporating Microbial Pilin-Based Nanowires into a Water-Stable Electronic Polymer Composite

**DOI:** 10.1101/2024.06.11.598525

**Authors:** Jayesh M. Sonawane, Eric Chia, Toshiyuki Ueki, Jesse Greener, Stephen S. Nonnenmann, Jun Yao, Derek R. Lovley

## Abstract

Electrically conductive protein nanowires (e-PNs), microbially produced from a pilin monomer, are a novel, sustainable electronic material that can be genetically tailored for specific functions. e-PNs, expressed with *Escherichia coli* grown on the biodiesel byproduct glycerol, and mixed with polyvinyl butyral yielded a transparent, electrically conductive water-stable composite.Composite conductivity was adjusted by modifying the e-PN concentration or incorporating e-PNs genetically tuned for different conductivities. Electronic devices in which composites were the sensor component differentially responded to dissolved ammonia over a wide concentration range (1µM-1M). Genetically modifying e-PNs to display an ammonia-binding peptide on their outer surface increased the sensor response to ammonia 10-fold. These results, coupled with the flexibility to design peptides for specific binding of diverse analytes, demonstrate that sustainably produced e-PNs offer the possibility of incorporating multiple sensor components, each specifically designed to detect different analytes with high sensitivity and selectivity, within one small sensor device.

## Introduction

Electrically conductive, microbially produced, pilin-based protein nanowires (e-PNs) show promise as a ‘green’ sustainably produced nanowire material for novel electronics applications [1,2]. More traditional nanowires comprised of silicon, carbon nanotubes, or metal require substantial energy for mining and/or synthesis, utilize toxic chemicals for synthesis, and the final products are toxic and/or not biodegradable [1-3]. In contrast, the microbes that assemble e-PNs from a pilin monomer protein can be grown on simple renewable carbon substrates (organic acids, alcohols, sugars) at moderate temperatures and the final product is robust in electronic applications, but non-toxic and biodegradable. Furthermore, the properties of e-PNs can be exquisitely genetically tailored to functionalize e-PNs in ways that more traditional nanowires cannot readily be modified. For example, pilin genes can be designed to encode a short peptide at the carboxyl end of the pilin protein that is displayed on the outer surface of the e-PNs [4]. Display of peptides that specifically bind analytes of interest can enhance the response of sensors in which the tailored e-PNs are the sensing component 100-fold [5]. The conductivity along the length of e-PNs has been tuned over 10-million-fold by modifying the aromatic amino acid content encoded in pilin genes [1,2,6,7].

Pilin-based e-PNs were first discovered in the electroactive microbe *Geobacter sulfurreducens* [8] in which they play an important role in the microbe’s electron transfer to extracellular Fe(III) oxide or other microbial species [9]. However, *G. sulfurreducens* is not a good candidate for mass production of e-PNs because it is a slow growing anaerobe that can produce other conductive filaments that can contaminate nanowire preparations [10]. Microbes capable of fast growth under aerobic conditions can heterologously express the *G. sulfurreducens* pilin gene to produce e-PNs with the same 3 nm diameter and conductance as the pilin-based e-PNs that emanate from *G. sulfurreducens* [11-13]. Expression of e-PNs with an *Escherichia coli* chassis coupled with simple e-PN harvesting strategies suggest that e-PN production is scalable for commercial applications [12]. *In vitro* synthesis of pilin-based e-PN mimics has been reported, but the high cost and toxic reagents required for pilin peptide synthesis and purification suggests that this approach may be more difficult to sustainably scale up than direct microbial production of e-PNs [1].

A diversity of electronic devices has been fabricated with thin-films of e-PNs. These include the ‘Air-gen’, which generates electricity from atmospheric humidity [14], memristor and neuromorphic memory devices [15,16], and sensors [5,17,18]. The e-PNs in these devices are robust, retaining full functionality over months to years [5,14,17-19]. e-PNs mixed with polyvinylalcohol (PVA) yielded an electrically conductive composite [20]. e-PN/PVA composite conductivities were comparable to composites of other nanowire materials and PVA with a relationship between e-PN density and e-PN/PVA composite conductivity consistent with percolation theory [20].

A limitation of e-PN thin films and e-PN/PVA composites is that water can dissolve PVA and wash away e-PNs, limiting applications in environmental and biomedical sensing. Polyvinyl butyral (PVB) is water insoluble and forms a robust, flexible, adhesive polymer [21]. Composites of silver nanowires and PVB yielded a conductive, transparent composite [22-24]. Here we report on the fabrication of water-stable, electrically conductive e-PN/PVB composites and demonstrate that the conductive and sensing properties of the composites can be tuned with genetic tailoring of the e-PN amino acid sequence.

## Results and Discussion

Mixtures of e-PNs in PVB at 2 weight percent e-PN were drop cast on a gold electrode array, forming a thin, uniform transparent film (Fig. 1a, b). Confocal scanning laser microscopy revealed that the e-PNs were uniformly distributed throughout the composite (Figure 1 c-f).

**Fig. 1.**
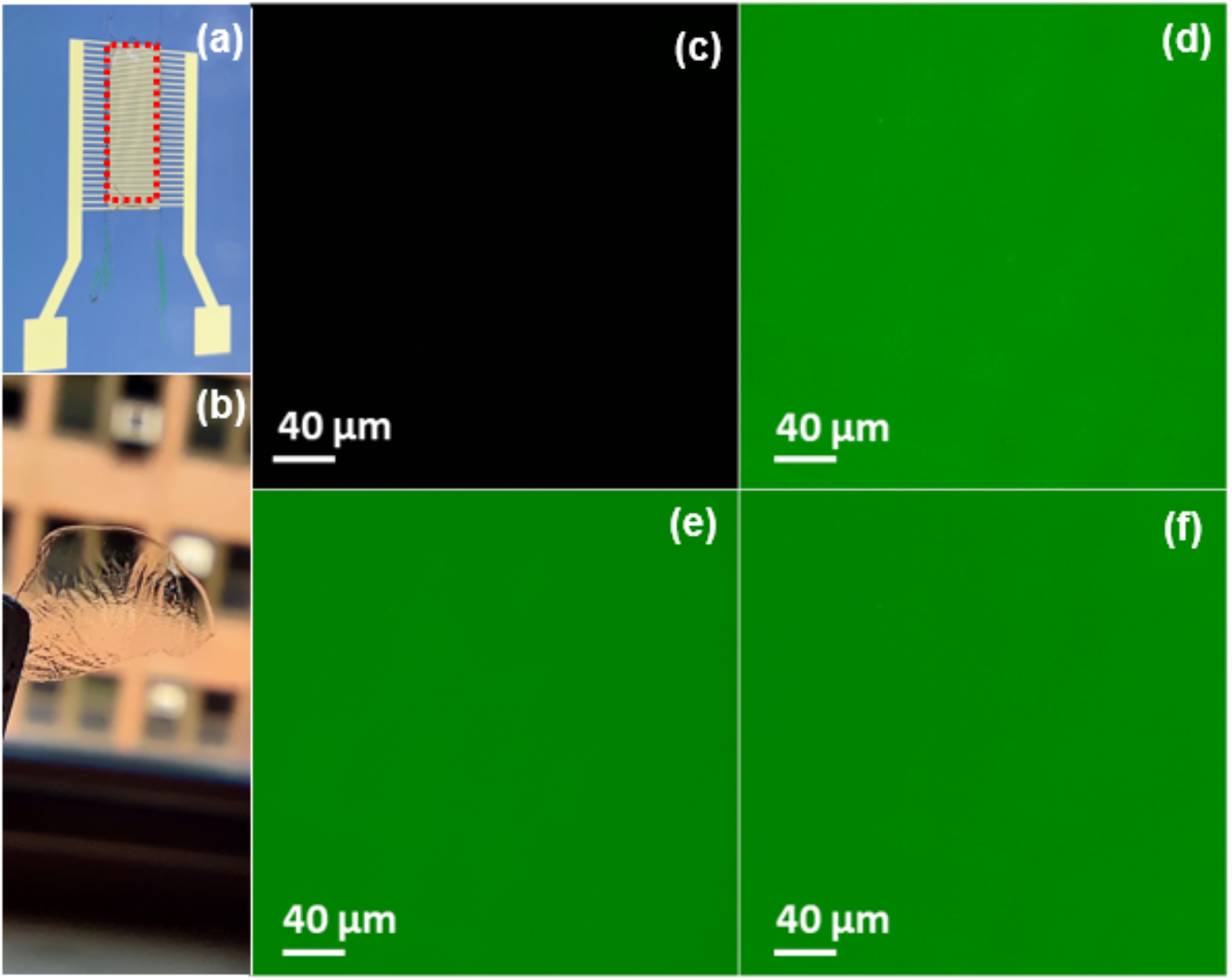
Appearance of e-PN/PVB composite at 2 weight percent e-PNs. (a) Composite drop cast on a gold electrode array. The position of the rectangular cured composite is highlighted with a red dotted outline. (b) Image demonstrating the transparency of a self-standing composite. (c-f) Confocal scanning laser microscopy imaging of e-PN distribution in the composite when e-PNs were fluorescently stained. (c) Control image in which e-PNs were not fluorescently stained. (d) Image of the average intensity of all slices along the Z axis. (e) First slice of Z scan (f) Last slice of the Z scan.

Evaluation of the current-voltage (I-V) response of the composites at different weight percent e-PN (Fig. 2a) revealed a nearly linear response (Fig. 2b), demonstrating ohmic contact between the composite and the electrodes. This result is consistent with previous reports on the near-ohmic response of individual e-PNs, e-PN thin-film networks, and composites of e-PNs with polyvinyl alcohol [6,8,20,25].

**Fig. 2.**
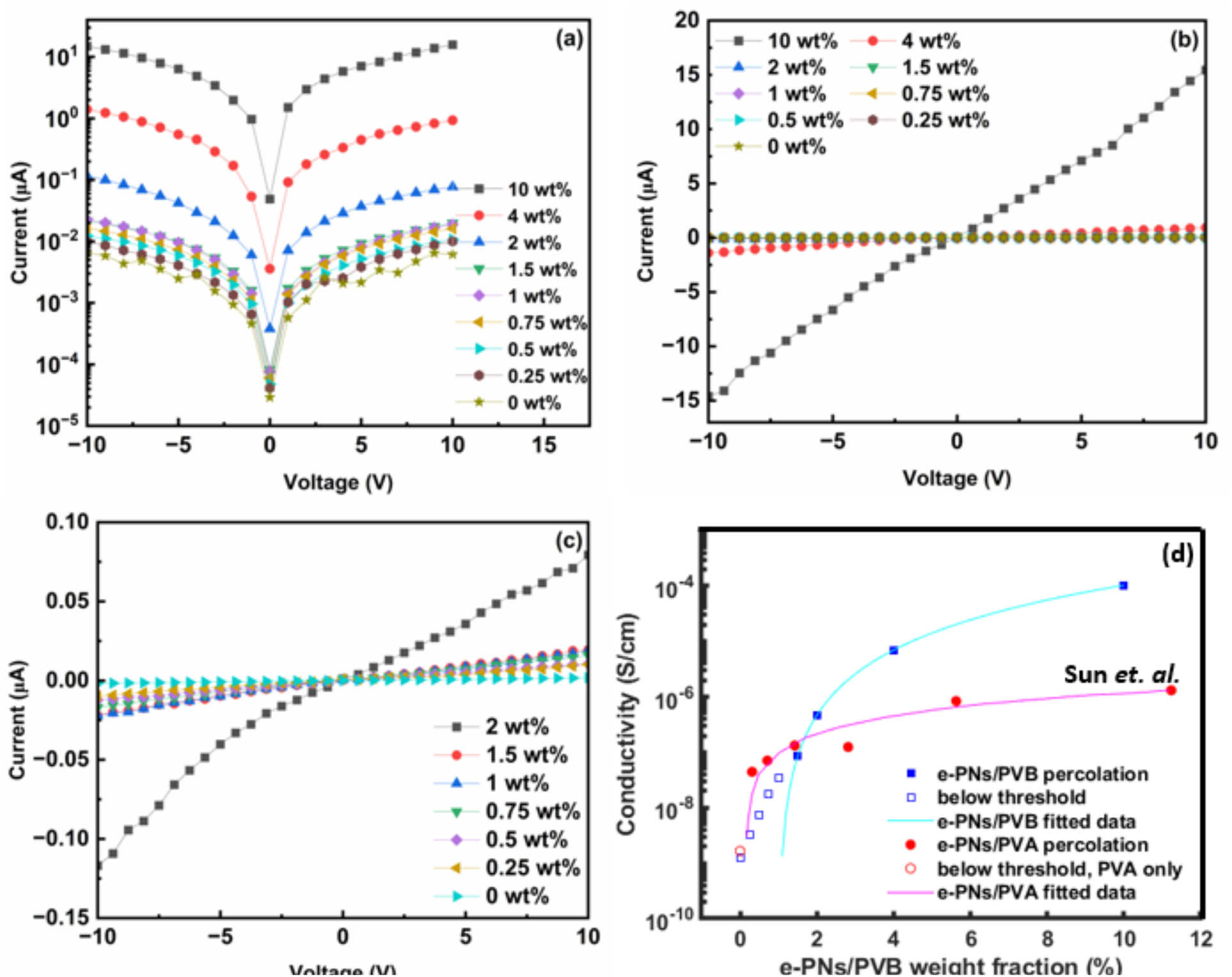
Impact of e-PN density on composite current response. (a) Current presented on logarithmic scale to more readily differentiate current response over the full range of weight percent e-PNs evaluated. (b, c) Linear presentation of data from panel (a). (d) Polymer conductivity versus e-PN weight percent from this study compared with the previously published data of Sun et al. [20] for e-PNs incorporated in a polyvinylalcohol composite.

As previously observed with e-PN/PVA composites [20], the conductivity of the e-PN/PVB composites increased with increasing weight percent e-PN (Fig. 2a-d), following classical percolation theory [26]:

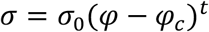

where σ is the electrical conductivity, σ_0_ is the pre-exponential constant, *φ* is the weight percentage of e-PN fillers, *φ*_*c*_ is the percolation threshold enabling connectivity and increased conductivity in the PVB matrix, and *t* is the critical exponent. Curve fitting yielded a *φ*_*c*_ of 1.01 weight percent (Fig. 2d). This threshold is higher than previously observed for e-PN/PVA composites (Fig. 2d), in which the observed percolation threshold matched well with what would theoretically be expected for randomly oriented, high aspect ratio nanowires [20]. One difference may be more e-PN bundling or other factors increasing heterogeneity within the PVB composites. Furthermore, PVB has a significantly lower dielectric constant than PVA [27,28] and the polymer dielectric constant can play a significant role in percolation [29-32], dictating the tunneling probability between conductive fillers in forming the percolative network [29,30]. At e-PN concentrations of 2 weight percent or more, conductivities of the e-PN/PVB composites were ca. 10-fold higher than previously reported conductivities for e-PN/PVA composites at comparable weight percent e-PN (Fig. 2d).

### Genetically tuning conductivity

The conductivity along the length of individual e-PNs and of e-PN thin films can be tuned by modifying the abundance of aromatic amino acids encoded in the *G. sulfurreducens* pilin gene [6,7,33]. Expressing an e-PN gene sequence in *E. coli* that was previously shown to yield e-PNs more conductive that the native sequence [7] yielded an e-PN/PVB composite more conductive than the composite fabricated with e-PNs harvested from the *E. coli* strain expressing the native e-PN gene sequence (Fig. 3). Expressing a gene that encoded e-PNs with a lower abundance of aromatic amino acids yielded e-PN/PVB composites with a lower conductivity.The relative difference in the conductivities of the composites were similar to differences in the conductivities of thin films of these e-PNs [25].

**Fig. 3.**
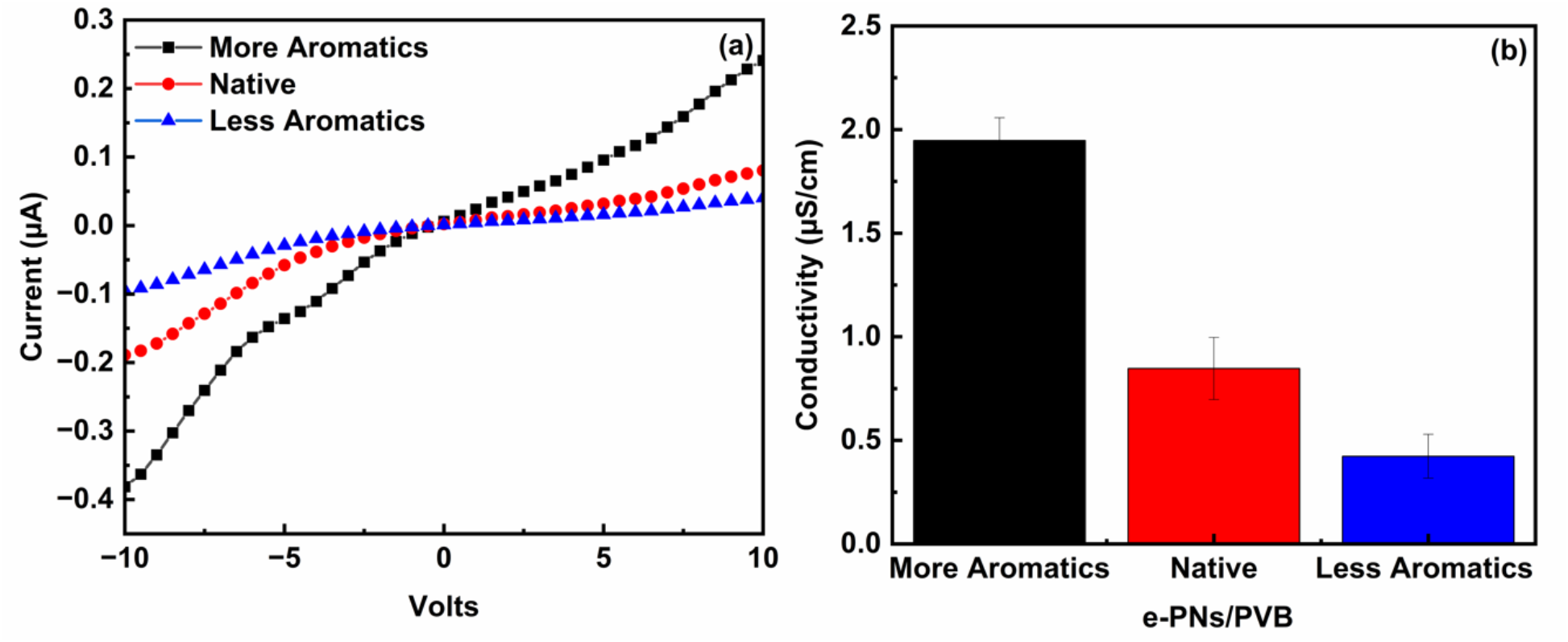
Impact on composite conductivity of modifying aromatic amino acid content encoded in e-PN pilin monomer genes expressed in *E. coli*. (a) Current versus voltage plots for composites (2 weight percent e-PN) fabricated with e-PNs with a higher or lower abundance of aromatic amino acids than the native e-PNs. (b) Conductivities calculated from data in panel a (mean + standard deviation for triplicate determinations.

### Tuning composite sensor sensitivity

Previous studies demonstrated that thin films of e-PNs harvested from *G. sulfurreducens* functioned as a sensing component for gaseous ammonia in an electronic device [17]. Genetic tailoring of the e-PN gene expressed in *E. coli* to display an ammonia-binding peptide (DLESFL) at the carboxyl end of the pilin monomer yielded e-PNs that conferred enhanced sensitivity for ammonia gas in thin-film devices [5]. To evaluate the potential for dissolved analyte sensing, e-PN/PVB composites were incorporated into an electronic sensing device held within a microfluidic system that provided a continuous flow of water over the composite. There was a rapid current response to the introduction of dissolved ammonia from the composites containing either e-PNs with the ammonia ligand or unmodified e-PNs (Figure 4 a-f). The current rapidly returned to baseline when the flow over the device was switched back to just water. The current response increased with increasing ammonia concentrations over the million-fold range of 1 µm to 1 M (Fig. 4 a-f). The sensitivity of the devices with composites of the e-PNs with the ammonia-binding peptide was ca. 10-fold greater than the response with the unmodified e-PN composites. This result is consistent with the previous finding that introducing the ammonia binding peptide enhanced the detection of gaseous ammonia [5]. However, the ammonia-binding ligand increased the response of e-PN thin films to gaseous ammonia ca. 100-fold over the response of the unmodified e-PNs.

**Fig. 4.**
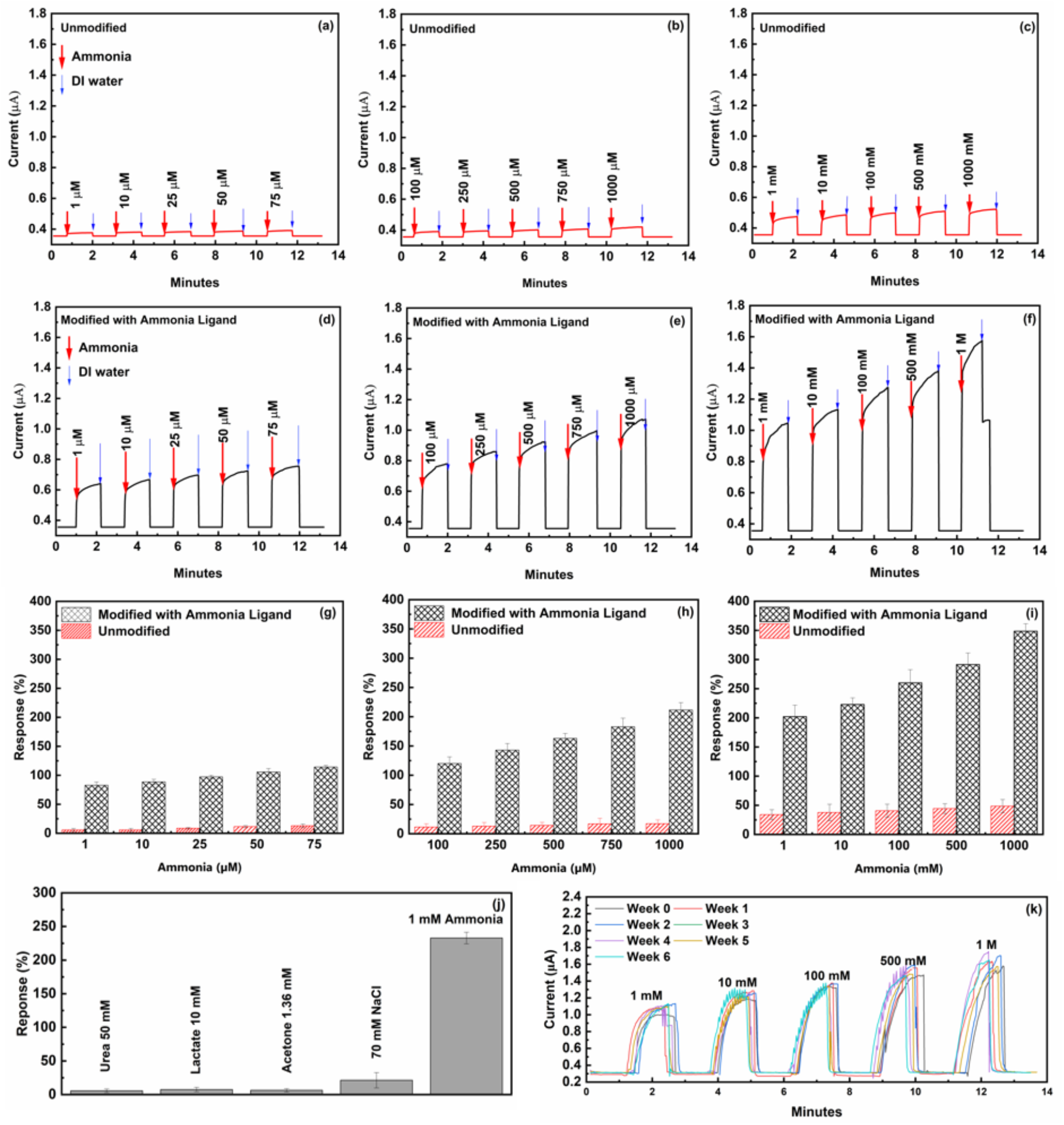
Sensor function of e-PN composites as the sensing component in an electronic device with a microfluidic aqueous input. (a-c) Current response to changes in the concentration of dissolved ammonia with unmodified e-PNs in the composite. (d-f) Current response to changes in the concentration of dissolved ammonia with e-PNs modified with an ammonia ligand. (g-i) Current response from panels a-f as a percentage of the baseline water-only current (mean + standard deviation of triplicate determinations). (j) Comparison of ammonia response to physiologically relevant concentrations of other sweat components. (k) Response of sensor fabricated with modified e-PNs to a range of ammonia concentrations from once-a-week testing over six weeks.

### Sensor Selectivity and Robustness

The current response of the devices fabricated with the e-PN/PVB composites with the ammonia ligand was specific. For example, ammonia concentrations in sweat can be diagnostic for various metabolic disorders, liver and kidney function, peptic ulcers, cancers, and metabolic exertion [34-38]. Therefore, the response of the composites with the e-PNs modified with the ammonia ligand to other common sweat constitutes (urea, lactate, acetate, sodium) were determined at physiologically relevant [39-42] sweat concentrations. The response to all these other sweat constituents was much lower than the response to 1 mM ammonia, a typical ammonia concentration in sweat (Figure 4 j).

In previous studies electronic devices fabricated with e-PN thin films did not lose function for as long as they were tested, with time frames ranging from several months [5,14,17,18] to three years [19]. e-PN/PVB composites fabricated with the e-PNs with the ammonia ligand did not lose function over a test period of six weeks (Figure 4 k). A one-way ANOVA statistical test for similarity yielded a p-value of 0.996.

## Conclusions

The results demonstrate that microbially produced e-PNs can be incorporated into a water-stable polymer to fabricate electrically conductive composites. This advance expands e-PN-based sensor analysis beyond volatiles and makes e-PNs available in a flexible, moldable form that is more stable than e-PN thin-film devices. The e-PNs were produced with microbes grown with glycerol, a waste product of biodiesel production. The e-PN/PVB composites were simply fabricated at room temperature, avoiding the use of toxic chemicals and high-energy inputs common with the synthesis of non-biological nanowires.

The properties of e-PNs are much more tunable than traditional nanowires comprised of metal, silicon, or carbon [1,2]. e-PN conductivity and the sensor response of the composites can be specifically customized with simple genetic modification of the amino acid content of the amino acids that comprise the e-PN monomer. The peptide ligands that bind analytes are integral components of the e-PNs, no additional chemistries are required for nanowire functionalization. Peptides that specifically interact with a diversity of environmentally and medically relevant analytes can readily be designed [43-51], providing the possibility of fabricating e-PNs to specifically detect a wide range of diverse compounds of medical and environmental interest.

Thus, e-PNs offer the option of incorporating highly specific and sensitive sensors for multiple dissolved analytes of interest in a single small device requiring little sample volume.

## Materials and Methods

### *E. coli* strains

*E. coli* strains expressing genes designed to produce the native *G. sulfurreducens* pilin protein monomer or a *G. sulfurreducens* pilin modified with the ammonia-binding peptide DLESFL [52] incorporated into the carboxyl terminus were previously described [5,12] and obtained from our laboratory culture collection.

A pilin gene, designated *aro5*, which encodes a pilin with lower aromatic amino acid abundance than the native *G. sulfurreducens* pilin and previously demonstrated to yield poorly conductive pili [6,25] was synthesized by Integrated DNA Technologies. The DNA sequence was:

tctcat

ATGGACAAGCAACGCGGTTTCACCCTTATCGAGCTGCTGATCGTCGTTGCGATCATC GGTATTCTCGCTGCAATTGCGATTCCGCAG**GC**CTCGGCG**GC**TCGTGTCAAGGCG**GC** CAACAGCGCGGCGTCAAGCGACTTGAGAAACCTGAAGACTGCTCTTGAGTCCGCA**G C**TGCTGATGATCAAACC**GC**TCCGCCCGAAAGTTAA
gagctcaga
(Nucleotides for the *E. coli* signal sequence are underlined. Nucleotides modified from the native pilin sequence to introduce alanine in place of aromatic amino acids are indicated in bold.) with the corresponding amino acid sequence of:

FTLIELLIVVAIIGILAAIAIPQ**A**SA**A**RVKA**A**NSAASSDLRNLKTALESA**A**ADDQT**A**PPES (Alanines that substitute for the aromatic amino acids are indicated in bold.)

The pilin gene from the microbe *G. metallireducens* yields e-PNs with higher conductivity than the native *G. sulfurreducens* pilin, presumably due to a higher abundance of aromatic amino acids in the pilin monomer [7]. The *G. metallireducens* pilin gene was produced from *G. metallireducens* genomic DNA template with PCR with the following primer pair: TCTCATATGGACAAGCAACGCGGTTTCACCCTCATCGAGCTGC and TCTGAGCTCTTAATTCGGATAAAATTGGTGTTC.

The amino acid sequence of the *G. metallireducens* pilin is:

FTLIELLIVVAIIGILAAIAIPQFAAYRQKAFNSAAESDLKNTKTNLESYYSEHQFYPN As previously described for other synthetic pilin genes introduced into *E*. coli [5,12], the signal sequence for the *E. coli* PpdD pilin was included in the pilin gene sequences to facilitate e-PN assembly of Aro5 pilins and *G. metallireducens* pilins. The synthesized DNA fragment containing *aro5* or PCR-amplified DNA fragment containing the *G. metallireducens* pilin gene was digested with NdeI and SacI and then cloned into the nanowire expression vector T4PAS/p24Ptac [12]. The resultant plasmids were transformed into *E. coli ΔfimAΔfliC*, a strain in which genes for FimA, the primary monomer for type I pili, and FliC, the structural flagellin of flagella, were deleted [5].

### Protein nanowire production

*E. coli* strains were routinely grown at aerobically 30 °C in LB medium. Cells were mass cultured for e-PN production as previously described [5,12]. Briefly, cells were grown for 48 h at 30 °C on agar-solidified M9 medium with 0.5 % glycerol as the energy source; 0.5 mM IPTG to induce pilin gene expression; and 50 μg/ml kanamycin. Cells were scrapped from the agar surface, suspended in M9 medium, and centrifuged to collect a cell pellet. The cell pellets were suspended in 50 ml ethanolamine buffer (150 mM, pH 10.5) for protein nanowire purification with ammonium sulfate precipitation, as previously described [5]. The purified e-PNs were dialyzed against deionized water several times, suspended in 2 ml of sterile water and stored at 4°C. The protein content of nanowire suspension was determined with a BCA protein assay kit (Thermo Pierce, USA).

### Preparation of e-PN/PVB Composites

To make composites, aliquots of the aqueous e-PN suspension were dried overnight in polypropylene tubes at 37°C. PVB was dissolved in anhydrous ethanol at concentrations ranging from 1 to 20 mg/ml to achieve the desired weight percent e-PNs. The dried e-PNs were resuspended in the PVB solution.

Interdigitated gold electrode arrays were fabricated as previously described [5,17]. Briefly, chromium and gold films (Cr/Au, 5/50 nm) were deposited on the surface of a silicon wafer with standard photolithography, deposition, and lift-off processes. For conductivity measurements, the electrodes were 8 mm x 5 mm with 10 interdigitated fingers, each separated by a 100-micron gap. For the microfluidic sensor assembly, electrodes were 10 mm x 4 mm with 20 interdigitated fingers, each separated by a 100-micron gap.

A 4 × 10 mm area was defined on the electrode arrays with scotch tape. 10 µl of the e-PN/PVB solution was drop cast inside the tape-bound region and the alcohol allowed to evaporate at 25 °C for 15 min to yield the composite film. The tape was then removed leaving a free-standing composite on the electrode arrays.

To examine the distribution of the e-PNs within the PVB matrix, the nanowires were first stained with fluorescein isothiocyanate (FITC) as previously described [53]. The stained e-PNs were dialyzed against water for three cycles to remove unbound stain. The aqueous e-PN suspension was dried at 37 °C, and then PVB solution was added to achieve a weight percent e-PNs of 2. An area of 4 x 10 mm was defined on a glass microscope slide with scotch tape and 10 µl of the e-PN/PVB suspension was drop cast inside the tape-bound region and the alcohol allowed to evaporate at 25 °C for 15 min to yield the composite film. The tape was then removed leaving a free-standing composite on the microscope slide. Images were acquired with a Nikon A1R-SIMe confocal scanning laser super-resolution microscope. The images were processed with the Fiji/ImageJ software package supplied by the National Institutes of Health.

### Microfluidic sensor device

To produce a microfluidic sensor device, a channel (width 2 mm; length 25 mm; depth 0.2 mm) with an inlet and an outlet port was fabricated in a block (45 mm x 35 mm) of polydimethylsiloxane (PDMS) with photolithography. The PDMS and silicon base of the electrode assembly were plasma treated to enhance the adhesion of the two hydrophobic materials. The channel was positioned over the e-PN/PVB composite on the electrode arrays described above and the assembly was incubated at 37 °C for four hours to promote adhesion. Water or water with dissolved analytes of interest was introduced at the inlet with a peristaltic pump at a flow rate of 5 ml per hour. To switch to a new analyte concentration, the valve on the pumping system was switched manually to introduce water into the channel at an elevated flow rate of 50 ml h^-1^ for 80 seconds to achieve baseline current to ensure that the microfluidic channel was flushed.

### Composite conductivity and sensor response

Current-voltage responses were determined with a semiconductor characterization system (Keithley 4200-SCS, Tektronix, Oregon, USA). The voltage range was -10V to 10V at a scan rate of 1mV/s. The relative humidity of the air was constant (25 ± 2%) throughout the analysis.

The conductivity was calculated from the current-voltage plots as:

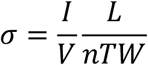

where *I* is the current between electrodes, *V* is the voltage across the two electrodes, *n* is the number of electrode pairs, *L* is the distance between the electrodes, *T* is the film thickness, as measured with a nano stylus profilometer (Bruker Dektak, Massachusetts, USA), and *W* is the cross-sectional width of the film [20].

The sensor responses were calculated using the following formula [5,54] :

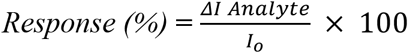

where *I*_0_ is the background current measured when DI water passed through the sensor, and ΔI represents the maximum change in current when the ammonia sample passed through the sensor:

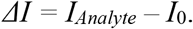

## Acknowledgments

J.Y. and D.R.L. acknowledge support from the National Science Foundation grant NSF-DMR-2027102. S.S.N. and D.R.L. acknowledge support from the National Science Foundation grant NSF-DMREF-1921839. Jayesh M. Sonawane was supported with a Fulbright-Nehru Fellowship.

## References

1. Lovley, D.R. and Yao, J. (2021) Intrinsically Conductive Microbial Nanowires for ‘Green’ Electronics with Novel Functions. Trends in Biotechnology 39, 940–952.

2. Guberman-Pfeffer, M.J. et al. (2024) Microbial nanowires for sustainable electronics. Nature Reviews Bioengineering 2, (in press).

3. Lovley, D.R. (2017) e-Biologics: Fabrication of sustainable electronics with ‘green’ biological materials. mBio 8, e00695–17.

4. Ueki, T. et al. (2019) Decorating the outer surface of microbially produced protein nanowires with peptides. ACS Synthetic Biology 8, 1809–1817.

5. Lekbach, Y. et al. (2023) Microbial nanowires with genetically modified peptide ligands to sustainably fabricate electronic sensing devices. Biosensors and Bioelectronics 226, 115147.

6. Adhikari, R.Y. et al. (2016) Conductivity of individual Geobacter pili. RSC Advances 6, 8354–8357.

7. Tan, Y. et al. (2017) Expressing the Geobacter metallireducens PilA in Geobacter sulfurreducens Yields Pili with Exceptional Conductivity. mBio 8, 10.1128/mbio.02203-02216. doi:10.1128/mbio.02203-16.

8. Reguera, G. et al. (2005) Extracellular electron transfer via microbial nanowires. Nature 435, 1098–1101. 10.1038/nature03661.

9. Lovley, D.R. and Holmes, D.E. (2022) Electromicrobiology: The ecophysiology of phylogenetically diverse electroactive microorganisms. Nature Reviews Microbiology 20, 5–19.

10. Liu, X. et al. (2021) Direct observation of electrically conductive pili emanating from Geobacter sulfurreducens. mBio 12, e02209–21.

11. Liu, X. et al. (2019) Biological synthesis of high-conductive pili in aerobic bacterium Pseudomonas aeruginosa. Appl. Microbiol. Biotechnol. 103, 1535–1544.

12. Ueki, T. et al. (2020) An Escherichia coli Chassis for Production of Electrically Conductive Protein Nanowires. ACS Synthetic Biology 9, 647–654. 10.1021/acssynbio.9b00506.

13. Szmuc, E. et al. (2023) Engineering Geobacter pili to produce metal:organic filaments. Biosensors and Bioelectronics 222, 114993.

14. Liu, X. et al. (2020) Power generation from ambient humidity using protein nanowires. Nature 578, 550–554.

15. Fu, T. et al. (2020) Bioinspired bio-voltage memristors. Nature Communications 11, 1861.

16. Fu, T. et al. (2021) Self-sustained green neuromorphic interfaces. Nature Communications 12, 3351.

17. Smith, A.F. et al. (2020) Bioelectronic protein nanowire sensors for ammonia detection. Nano Research 13, 1479–1484.

18. Liu, X. et al. (2020) Multifunctional protein nanowire humidity sensors for green wearable electronics. Advanced Electronic Materials 6, 2000721.

19. Liu, X. et al. (2023) Generic Air-Gen effect in nanoporous materials for sustainable energy harvesting from air humidity. Advanced Materials 2023, 2300748.

20. Sun, Y.-L. et al. (2018) Conductive Composite Materials Fabricated from Microbially Produced Protein Nanowires. Small 14, 1802624.

21. Kumar, P. et al. (2016) Polyvinyl butyral (PVB), versatile template for designing nanocomposite/composite materials: a review. Green Chemistry & Technology Letters 2, 185–194. 10.18510/gctl.2016.244.

22. Lee, D.J. et al. (2018) Light sintering of ultra-smooth and robust silver nanowire networks embedded in poly(vinyl-butyral) for flexible OLED. Scientific Reports 8, 14170.

23. Zhang, F. et al. (2020) Facile synthesis of Ag nanowires enhanced PVB for transparent conductive film. Journal of Materials Research and Technology 9, 14509–14516.

24. Li, G. et al. (2022) Ultrathin, flexible, conductive silver nanowires@polyvinyl alcohol composite film fabricated via the combination of air plasma treatment and thermal sintering for electromagnetic interference shielding. Materials Letters 325, 132814.

25. Vargas, M. et al. (2013) Aromatic Amino Acids Required for Pili Conductivity and Long-Range Extracellular Electron Transport in Geobacter sulfurreducens. mBio 4, 10.1128/mbio.00105-13.

26. Zare, Y. et al. (2022) Advancement of the Power-Law Model and Its Percolation Exponent for the Electrical Conductivity of a Graphene-Containing System as a Component in the Biosensing of Breast Cancer. Polymers 14, 3057.

27. Reddy, P.L. et al. (2019) Dielectric properties of polyvinyl alcohol (PVA) nanocomposites filled with green synthesized zinc sulphide (ZnS) nanoparticles. Journal of Materials Science: Materials in Electronics 30, 4676–4687.

28. Raghavendra, M. et al. (2022) Effect of CeO2 nanoparticles on dielectric properties of PVB/CeO2 polymer nanodielectrics: a positron lifetime study. Journal of Materials Science: Materials in Electronics 33, 1063–1077.

29. Sivadas, A. et al. (2022) Dielectric and Electrical Conductivity Studies of Carbon Nanotube-Polymer Composites. In Handbook of Carbon Nanotubes (Abraham, J.et al., eds), pp. 1209–1233, Springer International Publishing.

30. Zhang, L. et al. (2014) Revisiting the percolation phenomena in dielectric composites with conducting fillers. Applied Physics Letters 105, 042905.

31. Mazaheri, M. et al. (2022) Modeling of Effective Electrical Conductivity and Percolation Behavior in Conductive-Polymer Nanocomposites Reinforced with Spherical Carbon Black. Applied Composite Materials 29, 695–710.

32. Payandehpeyman, J. et al. (2024) Physics-Based Modeling and Experimental Study of Conductivity and Percolation Threshold in Carbon Black Polymer Nanocomposites. Applied Composite Materials 31, 127–147.

33. Tan, Y. et al. (2016) Synthetic Biological Protein Nanowires with High Conductivity. Small 12, 4481–4485.

34. Shalimar et al. (2019) Prognostic Role of Ammonia in Patients With Cirrhosis. Hepatology 70, 982–994.

35. Hu, C. et al. (2020) Serum ammonia is a strong prognostic factor for patients with acute-on-chronic liver failure. Sci Rep 10, 16970.

36. Silva, L.G. et al. (2022) Photoacoustic detection of ammonia exhaled by individuals with chronic kidney disease. Lasers Med Sci 37, 983–991.

37. Bai, C. et al. (2021) Urea as a By-Product of Ammonia Metabolism Can Be a Potential Serum Biomarker of Hepatocellular Carcinoma. Front Cell Dev Biol 9, 650748.

38. Kim, H. et al. (1990) The gastric juice urea and ammonia levels in patients with Campylobacter pylori. Am J Clin Pathol 94, 187–191.

39. Keller, R.W. et al. (2016) Urea transporters and sweat response to uremia. Physiol Rep 4, e12825.

40. Sakharov, D.A. et al. (2010) Relationship between lactate concentrations in active muscle sweat and whole blood. Bull Exp Biol Med 150, 83–85.

41. Li, J. et al. (2021) Aggregation kinetics of diesel soot nanoparticles in artificial and human sweat solutions: Effects of sweat constituents, pH, and temperature. J Hazard Mater 403, 123614.

42. Braconnier, P. et al. (2018) Sodium concentration of sweat correlates with dietary sodium intake. Journal of Hypertension 36, e170.

43. Wu, J. et al. (2011) Rapid Development of New Protein Biosensors Utilizing Peptides Obtained via Phage Display. PLOS ONE 6, e24948.

44. Liu, Q. et al. (2015) Peptide-based biosensors. Talanta 136, 114–127.

45. Vanova, V. et al. (2021) Peptide-based electrochemical biosensors utilized for protein detection. Biosensors and Bioelectronics 180, 113087.

46. Del Carlo, M. et al. (2016) In Silico Design of Short Peptides as Sensing Elements for Phenolic Compounds. ACS Sensors 1, 279–286.

47. Azmi, S. et al. (2015) Detection of Listeria monocytogenes with Short Peptide Fragments from Class IIa Bacteriocins as Recognition Elements. ACS Combinatorial Science 17, 156–163.

48. Kim, D.T.H. et al. (2018) Development of a novel peptide aptamer-based immunoassay to detect Zika virus in serum and urine. Theranostics 8, 3629–3642.

49. Tertis, M. et al. (2021) Electrochemical Peptide-Based Sensors for Foodborne Pathogens Detection. Molecules 26, 3200.

50. Pardoux, E. et al. (2019) Antimicrobial peptide arrays for wide spectrum sensing of pathogenic bacteria. Talanta 203, 322–327.

51. Wasilewski, T. et al. (2022) Olfactory receptor-based biosensors as potential future tools in medical diagnosis. Trends in Analytical Chemistry 150, 116599.

52. McAlpine, M.C. et al. (2008) Peptide− nanowire hybrid materials for selective sensing of small molecules. Journal of the American Chemical society 130, 9583–9589.

53. Chaganti, L.K. et al. (2018) An efficient method for FITC labelling of proteins using tandem affinity purification. Biosci Rep 38, BSR20181764.

54. Jha, R.K. et al. (2018) Ammonia vapour sensing properties of in situ polymerized conducting PANI-nanofiber/WS2 nanosheet composites. New Journal of Chemistry 42, 735–745.

